# *Peptonella octanoica* gen. nov., sp. nov., a new medium-chain carboxylate-producing bacterium, and the reclassification of *Eubacterium pyruvativorans* as *Peptonella pyruvativorans* comb. nov

**DOI:** 10.64898/2026.08.23.746564

**Authors:** Dinesh Kumar Nallasamy, Blake G Lindner, Christopher E Lawson

## Abstract

A strictly anaerobic bacterial strain, F2^T^, was isolated from an anaerobic bioreactor fermenting source-separated organic waste. Cells of strain F2^T^ are non-spore-forming, rod-shaped (1.5–2.5 × 0.27–0.33 µm), and Gram-negative, although they possess a monoderm cell wall architecture. The strain grew at 37 °C within a pH range of 5 to 8 and produced short-, branched-, and medium-chain carboxylates as well as ammonium, H_2_ and CO_2_, with acetate and propanoate produced or consumed depending on fermentation conditions. The genome consists of a single 2.4 Mbp chromosome with a G+C content of 50.2% and 2,131 predicted genes. Phylogenetic analysis of the 16S rRNA gene against other isolates revealed that strain F2^T^ is most similar to *Eubacterium pyruvativorans* I-6^T^ (92.06% 16S rRNA identity). Based on further phenotypic, genomic, and phylogenetic analysis, strain F2^T^ represents a novel genus and species within the family *Anaerovoracaceae* with the proposed name *Peptonella octanoica* gen. nov. sp. nov. The type strain is F2^T^ (strain accession pending). As a member of this same genus-level clade, we propose reclassifying *Eubacterium pyruvativorans* as *Peptonella pyruvativorans* comb. nov. These findings disambiguate *Peptonella spp.* from the phylogenetically distant and phenotypically distinct *Eubacterium limosum* ATCC 8486^T^.

## Main Text

Medium-chain carboxylic acids (MCCAs) constitute a class of chemicals comprising 6 to 12 carbon atoms that have various applications as feed additives, antimicrobials, and cosmetics [1–4]. MCCAs are synthesized via an anaerobic bioprocess termed microbial chain elongation, where acyl-CoA units are extended by two carbons per reverse β-oxidation (rBOX) cycle. Chain elongation is facilitated by electron donors such as ethanol or lactate, and has emerged as a promising biotechnological platform for transforming organic waste streams into these valuable products [2, 3]. Interest in this platform has increased significantly over the past decade, supported by long-term bioreactor studies that demonstrate consistent MCCA production from diverse organic feedstocks [1, 5].

Most reports on MCCA production to date have used open-culture microbial communities as biocatalysts, including commercial efforts [6]. While open cultures are effective for converting waste feedstocks, their use complicates understanding the underlying microbiology and microbial ecology [6, 7]. Consequently, key metabolic principles governing MCCA production metrics, such as product yield and chain length, as well as microbial interactions governing carbon flux remain only partially understood [6, 8].

The collection of formally characterized CEBs is predominantly derived from environmental or host-associated sources and exhibits minimal resemblance to chain elongators observed in bioreactors processing complex organic waste [5, 9]. *Clostridium kluyveri*, recognized as the model CEB and the most extensively studied member of this functional group, was initially isolated from canal mud in the late 1930s [10, 11]. Other chain-elongating isolates include *Megasphaera hexanoica*, *Megasphaera elsdenii*, and *Eubacterium pyruvativorans*, all of which were isolated from the rumen of cattle or sheep [5, 12–14]. Several additional isolates, including *Megasphaera cerevisiae* and *Pseudoramibacter alactolyticus,* were incidentally recovered due to their roles in beer spoilage or their association with human oral diseases, rather than through targeted isolation campaigns focused on MCCA-producing bioreactor communities [6]. *Caproiciproducens galactitolivorans* and *Caproiciproducens sp.* 7D4C2 are among the few CEB isolated from bioreactor enrichments fed sugars cultivated at mildly acidic pH conditions [5, 9]. Yet, the isolation record is far from complete; metagenomic surveys have identified a broad distribution of putative chain elongators within the families *Anaerovoracaceae*, *Eubacteriaceae*, *Lachnospiraceae*, and *Oscillospiraceae* that have yet to be isolated [6, 7]. Consequently, currently available isolates do not comprehensively represent the expected functional diversity of CEB inhabiting waste-processing bioreactor environments.

Here, we present the isolation and characterization of strain F2^T^, a novel CEB obtained from an anaerobic bioreactor fed with source-separated organics (SSO) and operated for MCCA production, as reported by Dyussekenova and Parmar et al. [15]. Strain F2^T^ generates n-octanoate (C8) as a major product in modified DSM 104 media supplemented with lactate, distinguishing it from most characterized CEB, which predominantly produce n-hexanoate (C6). Based on phenotypic and phylogenomic analysis, we propose that strain F2^T^ constitutes the type strain of a new genus and species, *Peptonella octanoica* gen. nov., sp. nov., and that *Eubacterium pyruvativorans* [Wallace et al. 2003] should be reclassified as *Peptonella pyruvativorans* comb. nov.

## Isolation and cultivation

An anaerobic bioreactor producing MCCAs from organic waste was used as the source for isolating novel CEBs. The reactor was fed SSO containing lactate and short-chain carboxylates (SCCAs) that varied by season and collection facility, and was operated continuously at a solid retention time of 8-12 days, temperature of 37 °C, and pH 5 [1]. Samples for the isolation campaign were obtained from the reactor before the installation of the in-line MCCA extraction setup. The sample (an aliquot from the reactor) was serially diluted inside an anaerobic chamber. 100 µL of 10^-2^ and 10^-3^ dilutions were spread-plated on different synthetic media agar plates (pH 6) that were pre-reduced in the anaerobic chamber, rendering them oxygen-free. The plates were placed in an anaerobic jar (BD GasPak jars) and accompanied by a BD Anaerobe Container System Sachet containing an indicator and a BD CO_2_ indicator strip. This jar was then incubated at 37 °C for 7 days. Colonies with distinct morphologies were picked using a sterile toothpick (inside an anaerobic chamber), transferred to the respective liquid media, and incubated anaerobically for 3–5 days. Glycerol stocks were prepared from the liquid culture, and a portion of the culture was passaged into the respective media in a 96-well plate to screen metabolites. Several anaerobic isolates were obtained, among which strain F2^T^ was selected for detailed characterization based on its distinctive MCCA production profile.

Strain F2^T^ was isolated under anaerobic conditions using the media (modified DSM medium 104 – mDSM104) containing 10 g/L sodium lactate, 5 g/L tryptose, 5 g/L peptone, 10 g/L yeast extract, 5 g/L beef extract, 2 g/L K_2_HPO_4_, 1 mL/L Tween 20, 40 mL/L salt solution, 1 mL/L of 0.1% (w/v) resazurin, 0.2 mL/L of vitamin K_1_ solution, 10 mL/L of haemin solution, and 0.5 g/L of cysteine-HCL. The salt solution contained: 0.25 g/L CaCl_2_.2H_2_O, 0.5 g/L MgSO_4_.7H_2_O, 1 g/L K_2_HPO_4_, 1 g/L KH_2_PO_4_, 10 g/L NaHCO_3_ and 2 g/L NaCl. The vitamin K1 solution was prepared by dissolving 0.1 mL of vitamin K1 in 20 mL of 95% ethanol and filter-sterilizing. The haemin solution was prepared by dissolving 50 mg of haemin in 1 mL of 1N NaOH, adjusting the volume to 100 mL with deionized water, and then filter-sterilizing. 15 g/L agar was used to prepare solid media. The pH of the media was adjusted to 6.0±0.2 and sterilized by autoclaving at 121 °C for 30 minutes. Vitamin K_1_ and haemin solutions were added to the media after autoclaving, and the plates were kept in the anaerobic chamber for 2 days to remove oxygen.

## Morphology and physiology

Cell and colony morphology were evaluated by cultivating strain F2^T^ on mDSM104 medium supplemented with lactate at pH 6, followed by incubation at 37 °C for 5 days. Gram staining of strain F2^T^ was conducted in parallel with a Gram-positive bacterium (*Clostridium kluyveri* DSM 555) and a Gram-negative bacterium (*Escherichia coli* BL-21), utilizing a Gram stain kit (77730-1KT-F, Sigma-Aldrich) in accordance with the manufacturer’s instructions. The stained cells were examined using an optical microscope (Olympus BX51 – Bright Field, 100x magnification). Additionally, Transmission Electron Microscopy (TEM) analysis of strain F2^T^ was performed. The cell pellet was high-pressure frozen using a Leica EM ICE and then freeze-substituted in 2% osmium tetroxide in acetone at –90 °C. The sample was warmed to –30 °C, washed with acetone, and then infiltrated with Spurr resin. Fresh resin changes were made at room temperature, and blocks were cured in the oven at 60 °C overnight. Ultrathin sections were cut using a Leica UC7 ultramicrotome, and the grids were post-stained with uranyl acetate and lead citrate prior to viewing using a Hitachi HT7800 TEM. Endospore staining was carried out as previously described [16]. *Clostridium sporogenes* CE3 (JCAOOO000000000), an isolate from the same reactor, was used as a positive control for endospore staining.

Cells of Strain F2^T^ are rod-shaped and Gram-negative, as shown in Figure 1a. The shape and size (1.5–2.5 × 0.27–0.33 µm) of the cells were confirmed by TEM. Moreover, TEM images showed a typical Gram-positive cell-wall ultrastructure (Fig. 1b, c). In Fig. 1c, the cytoplasmic membrane, cell wall, and an additional surface layer are indicated by arrows. The cell wall should be 30–50 nm thick to stain Gram-positive, and a minimum cell wall thickness is required to retain the Gram-stain complex during decolorization [17]. Several other bacteria have been described that stain Gram-negative yet possess a Gram-positive cell wall structure, including *Caproicibacter fermentans* strain EA1^T^ (a chain-elongating bacterium) [9]. The cells of strain F2^T^ are non-sporulating, as indicated by endospore staining compared with the positive control. Also, no spores were observed in the TEM analysis. The colonies of strain F2^T^ on agar plates are small, white, circular, raised, and entire in margin.

**Fig. 1.**
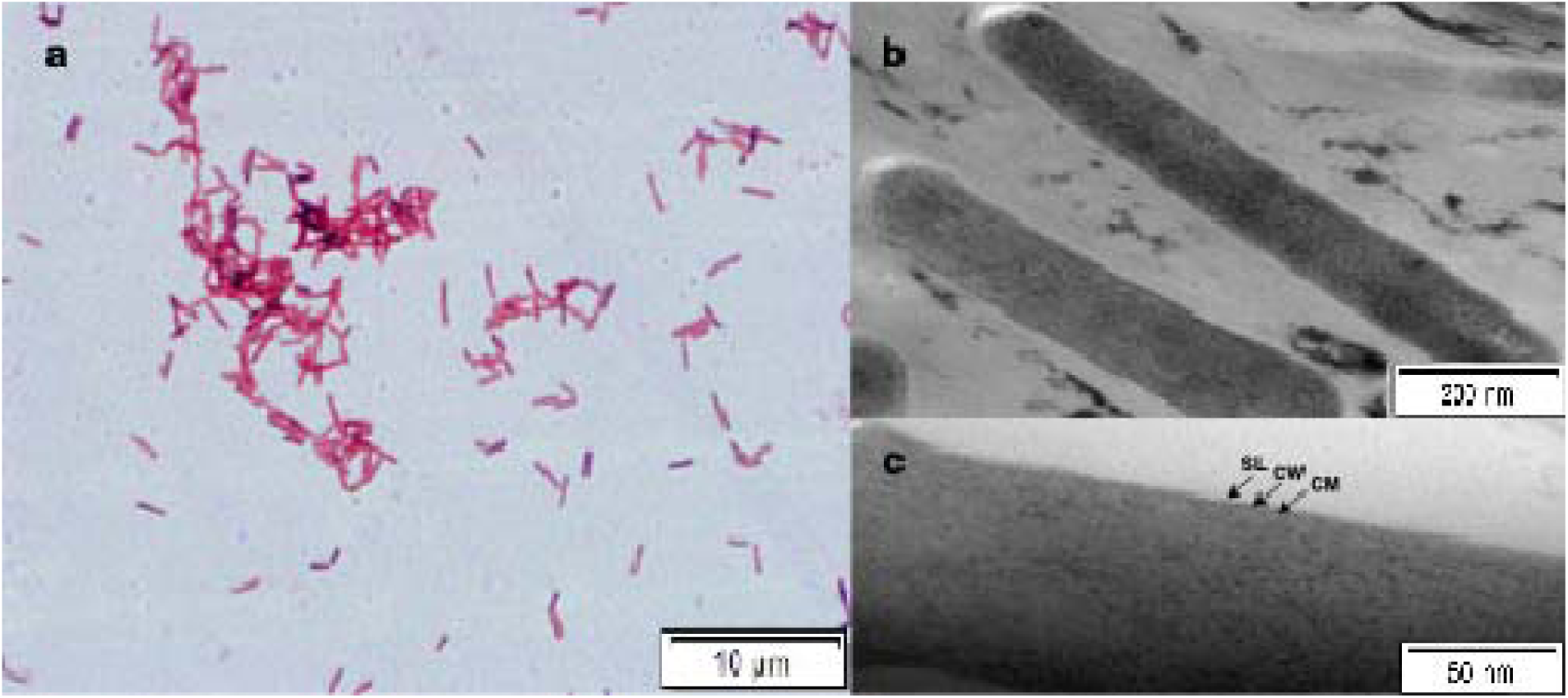
Microscopic images of strain F2^T^ cells. Gram-stained cells under an optical microscope (a) 100x fold magnification. TEM images using (b) 30000X fold magnification, (c) 120000X fold magnification. CM – cytoplasmic membrane; CW – cell wall; SL – surface layer.

Growth of strain F2^T^ at different pH was examined with initial values adjusted to 4, 5, 6, 7, and 8. Growth was observed across all values except at pH 4, with optimal growth between 5 and 6. Subsequently, experiments to determine the substrate preference of strain F2^T^ were performed at 37°C and pH 6. No difference in growth curves was observed when amending the media with glucose, galactose, xylose, mannose, lactose, galactitol, mannitol, glycerol, trehalose, arabinose, or cellobiose. However, a slight increase in biomass (<2 fold) was observed with the addition of fructose, sucrose, or ethanol. Lastly, the addition of lactate supported substantially elevated growth (>2 fold).

The products of strain F2^T^ cultivated in mDSM104 medium with added lactate were analyzed by GC-MS and HPLC. Lactate concentration was quantified using a high-performance liquid chromatograph (UltiMate 3000 HPLC system, Thermo Scientific, USA) with an Aminex HPX-87H column (Bio-Rad, USA). Samples were centrifuged at 10,000 rcf for 10 minutes, diluted with HPLC-grade water, and filtered through a 0.22-μm filter prior to analysis. The mobile phase was 5 mM sulfuric acid at a constant flow rate of 0.6 mL/min. The column temperature was set to 50°C, and UV detection was set to 214 nm. Refractive index and UV detection were used to identify and quantify lactate. Carboxylate products were quantified using an Agilent 8890 gas chromatograph (GC) equipped with tandem mass spectrometry (7000D Triple Quadrupole, Agilent, USA). Filtered samples were diluted with HPLC-grade water and formic acid to a final concentration of 0.1 M. The run time was 16 minutes, with a temperature gradient set to: 80°C for 1 minute, a 20°C min-1 ramp to 200°C, a 5°C min-1 ramp to 210°C, and a 20°C min-1 ramp to 250°C, then held for 5 minutes. Dynamic multiple reaction monitoring (dMRM) was used for mass spectrometry. Electron ionization was used to fragment ions of interest. Precursor and product ions for each compound were selected using Agilent software (Optimizer). Helium was used as the carrier gas at a flow rate of 1 mL/min.

Strain F2^T^ produced MCCAs, predominantly n-octanoate (16.3 mM), in mDSM104 supplemented with lactate (initial concentration of 57 mM). In addition to n-octanoate, the strain produced acetate (20.3 mM), n-butanoate (1.5 mM), and n-hexanoate (8.3 mM), with trace amounts of propanoate, isobutanoate, isopentanoate, pentanoate, isohexanoate, 5-methylhexanoate, and n-heptanoate. Elevated headspace pressure was observed during cultivation, consistent with the production of gases such as CO_2_ and H_2_ during chain elongation fermentation [9, 13]. Furthermore, the strain produced ammonium through deamination of amino acids derived from background media components such as peptone and yeast extract.

## 16S rRNA gene sequence analysis

Genomic DNA from strain F2^T^ was extracted using Qiagen’s QIAamp DNA Kits. The 16S rRNA gene was amplified by PCR with the universal primers 27F and 1492R. The amplicon was then sequenced by Sanger sequencing (TCAG facility at SickKids, Toronto), yielding a 1,241-base-pair sequence that has been deposited in GenBank under accession number PV739030. BLAST searches against the NCBI nucleotide database indicated that the closest validly described relatives of strain F2^T^ are *Eubacterium pyruvativorans* I-6^T^ (92.06% similarity), *Aminicella lysinilytica* WN037^T^ (90.12% similarity), and *Hornefia butyriciproducens* WCA-MUC-591-APC-3H^T^ (88.54% similarity). All comparative values are significantly below the 98.7% threshold generally used to define species boundaries based on 16S rRNA gene sequence identity [25], thereby suggesting that strain F2^T^ constitutes a novel species. Additionally, similarity values below 94.5% when compared to *A. lysinilytica* and *H. butyriciproducens* are consistent with a taxonomic distinction at the genus level from those related taxa [26].

The 16S rRNA gene sequences of strain F2^T^, *Eubacterium pyruvativorans* I-6^T^, and closely related type strains were aligned with MUSCLE v5.2 [18]. Evolutionary distances were computed using Kimura’s two-parameter model [19], and a neighbour-joining (NJ) phylogenetic tree was constructed following Saitou and Nei [20] with 1,000 bootstrap replicates [21]; only bootstrap values ≥ 10% are shown at internal nodes. An independent maximum-likelihood (ML) tree was reconstructed with FastTree v2.1.11 [22] under the GTR+Γ model. The NJ tree, Kimura two-parameter distances, and bootstrap consensus were computed using DistanceCalculator, DistanceTreeConstructor, and get_support modules of the Biopython package v1.84. *Eubacterium limosum* ATCC 8486^T^ served as the outgroup.

Strain F2^T^ and *Eubacterium pyruvativorans* I-6^T^ formed a distinct clade with 61% bootstrap support, nested within a broader assemblage that includes all *Aminicella species* recognized in the GTDB taxonomy [23] (Fig. 2). *Aminicella lysinilytica* DSM 28287^T^, the type strain of the genus *Aminicella*, was the immediate sister lineage to the F2^T^/I-6^T^ clade, with 45% bootstrap support. The entire ingroup, comprising all *Aminicella* species, *Mogibacterium pumilum* ATCC 700696^T^, and the two focal strains, received 100% bootstrap support, whereas *Eubacterium limosum* ATCC 8486^T^ constituted a deeply divergent outgroup, consistent with its classification within a distinct family (*Eubacteriaceae*).

**Fig. 2.**
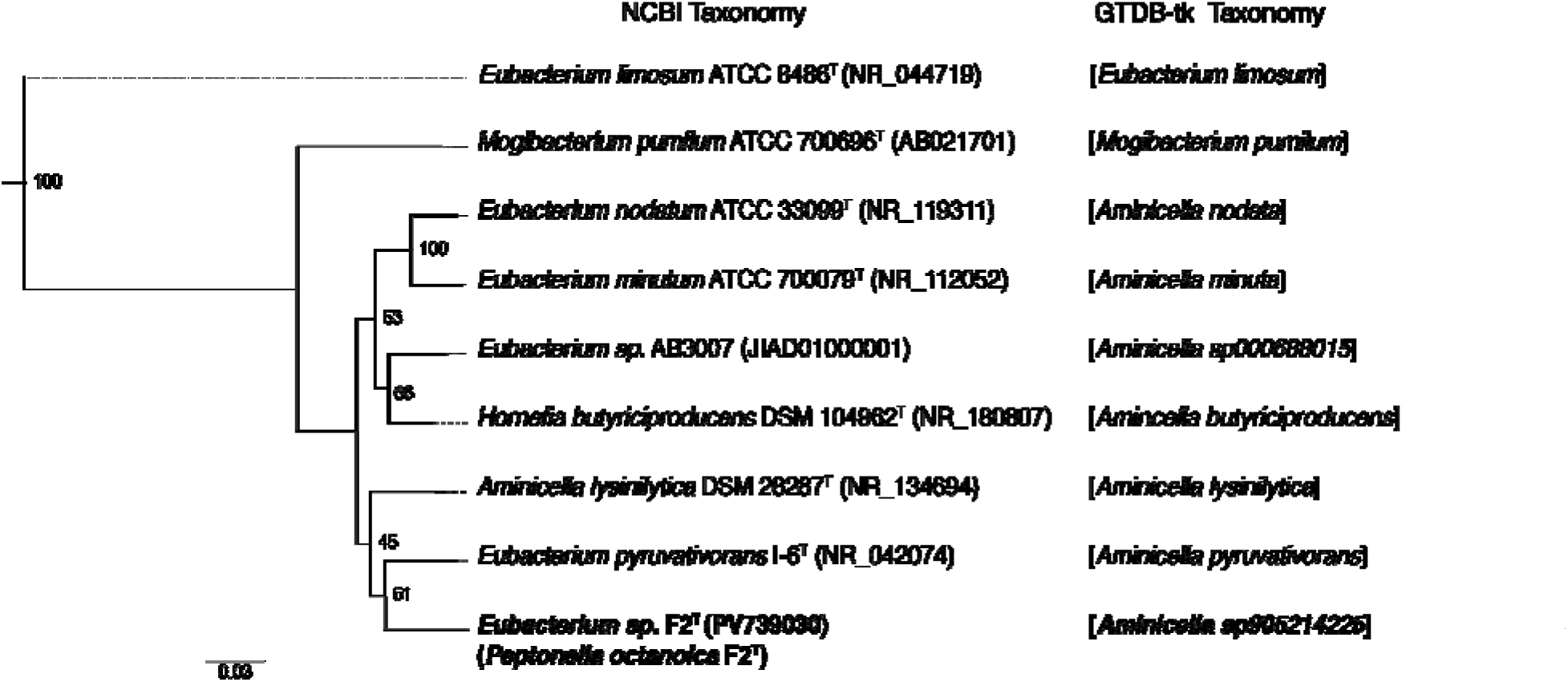
Neighbour-joining tree based on 16S rRNA gene sequences showing the phylogenetic position of *Peptonella octanoica* strain F2^T^ and related bacterial strains with nomenclatures from the NCBI and GTDB-tk databases. The phylogenetic tree was rooted to *Eubacterium limosum* ATCC 8486^T^. GenBank accession numbers of the 16S rRNA gene sequences are given in parentheses (for strain AB3007, the accession number is the whole-genome accession from which the 16S rRNA gene sequence was obtained). The maximum-likelihood tree was fully congruent with the NJ tree. Bootstrap values (based on 1000 replications) greater than or equal to 10% are shown as percentages at each node. Bar, 0.03 substitutions per nucleotide position.

These findings indicate that strain F2^T^ and *Eubacterium pyruvativorans* I-6^T^ belong to the family *Anaerovoracaceae* and form a distinct lineage within the *Aminicella* group. This lineage distinguishes them from *Eubacterium sensu stricto*, which is represented by the type species, *E. limosum* [23]. We observed relatively modest bootstrap support (61%) for the F2^T^/I-6^T^ clade with 16S rRNA analysis. Given the limited resolution of single-gene analysis, our investigation continued with whole-genome sequences for phylogenetic analysis.

## Genome sequencing and features

The genome of strain F2^T^ was sequenced using an Oxford Nanopore Technologies (ONT) platform (Plasmidsaurus, USA), resulting in 145,722 raw reads. These reads were assembled with Flye V2.9.1 [25], employing parameters optimized for high-quality ONT reads. The assembly yielded a single contig of 2,425,940 bp. The genome has a G+C content of 50.2 mol% and contains 2,131 genes, including 2,026 predicted protein-coding sequences, 5 rRNA genes (5X5S, 5X16S, and 5X23S), and 57 tRNAs. Genome completeness was assessed as 99.29% complete with 0% contamination using CheckM v1.2.2 [26].

The genome sequence of strain F2^T^ is available in both NCBI’s GenBank (GCA_046244355.1) and RefSeq databases (GCF_046244355.1). Analysis of the genome sequence of strain F2^T^ confirmed the presence of genes responsible for lactate oxidation and the gene cluster encoding the rBOX pathway, which is responsible for chain elongation.

The genome of *Eubacterium pyruvativorans* KHGC13 (DSM 118842) was sequenced on an ONT MinION (R10.4.1) with libraries prepared using the Rapid Barcoding Kit V14. The reads were assembled with Autocycler v0.5.2 and Canu v2.3, yielding a single 2,386,435-bp contig. The genome has a G+C content of 54.2 mol% and contains 2,230 genes, including 2,059 protein-coding sequences, 5 rRNA genes (5X5S, 5X16S, and 5X23S), and 60 tRNAs. The genome sequence is available in both NCBI’s GenBank (GCA_059427965.1) and RefSeq (GCF_059427965.1) databases. This strain also encodes the gene cluster for the rBOX pathway and lactate oxidation.

Both genomes contain a complete gene cluster encoding the Rnf complex, which functions as a ferredoxin: NAD^+^ oxidoreductase [EC 1.18.1.3] and oxidizes reduced ferredoxin (Fd2-) to generate NADH. ATP synthesis is driven by an ion gradient established as H^+^ or Na^+^ ions traverse the cytoplasmic membrane via ATPase. These genes are also found in other CEBs and perform a similar function [9].

## Digital DNA-DNA hybridization and species delineation

In silico DNA-DNA hybridization (dDDH) values were calculated using the Genome-to-Genome Distance Calculator (GGDC) 4.0 via the Type (Strain) Genome Server (TYGS) to determine the species status of strain F2^T^ and to confirm the species identity of *Eubacterium pyruvativorans* KHGC13. All values reported here use Formula 2 (the sum of all identities found in HSPs divided by the overall HSP length), the recommended formula for draft genomes, as it is independent of genome length [27].

The genomic analysis of strain F2^T^ compared with the 20 nearest type-strain genomes from the TYGS database identified it as a potential novel species (see Table S1). Notably, no type-strain genome exceeded the 70% dDDH species delineation threshold [27]. The highest dDDH value for strain F2 was 27.7% against *Eubacterium nodatum* ATCC 33099^T^ (*Aminicella nodata* in GTDB), followed by 26.5% against *Aminicella lysinilytica* DSM 28287^T^ (95% CI: 24.2–29.0%; probability of DDH ≥70%: 0.02%; G+C difference: 4.43%), and 21.6% against *Eubacterium pyruvativorans* I-6^T^ (95% CI: 19.3– 24.0%; probability of DDH ≥70%: 0%; G+C difference: 4.67%). All observed values remain substantially below the 70% species demarcation threshold, thereby confirming that strain F2 constitutes a novel species.

The pairwise GGDC analysis of *Eubacterium pyruvativorans* KHGC13 against the type strain I-6^T^ resulted in a dDDH of 84.9% (95% CI: 82.2–87.3%, probability of DDH ≥70%: 93.68%; G+C difference: 0.62%), confirming that strain KHGC13 is conspecific with the type strain of *Eubacterium pyruvativorans*. The dDDH between strain KHGC13 and strain F2^T^ was 23.0% (95% CI: 20.7–25.4%; probability of DDH ≥70%: 0%; G+C difference: 4.05%), and between strain KHGC13 and *Aminicella lysinilytica* DSM 28287^T^ was 24.9% (95% CI: 22.6–27.4%), with a probability of DDH ≥70%: 0.01%; G+C difference: 8.48%), both indicating species-level distinctness of the strains from all referenced taxa. The genome sequence of *Eubacterium pyruvativorans* KHGC13 was used for the preceding analysis.

## Genome-based phylogenetic analysis

To obtain a higher-resolution phylogenetic framework for strain F2^T^, a genome-based phylogenetic tree was inferred from a concatenated alignment of 120 conserved bacterial marker genes identified with GTDB-Tk v2.7.2 [28] using the de novo workflow against the GTDB release r232 reference database [23]. The concatenated alignment was trimmed from 41,084 to 5,010 amino acid positions after canonical masking implemented in the GTDB-Tk v2.7.2 and subsequently pruned to include the nine taxa of interest: Aminicella sp905214225 (*Peptonella octanoica* F2^T^), *Aminicella* (*Peptonella*) *pyruvativorans* KHGC13, *Aminicella lysinilytica* DSM 28287^T^, *Aminicella nodata* ATCC 33099^T^, *Aminicella minuta* ATCC 700079^T^, *Aminicella butyriciproducens* DSM 104962^T^, *Aminicella sp000688015* AB3007, *Mogibacterium pumilum* ATCC 700696^T^, and *Eubacterium limosum* ATCC 8486^T^ as the outgroup. Maximum-likelihood inference was performed using IQ-TREE v2.3.6 [29] with automatic model selection by ModelFinder [30] and 1,000 ultrafast bootstrap replicates [31].

ModelFinder identified Q.insect+F+G4 as the most appropriate substitution model based on the Bayesian information criterion (BIC). The phylogenetic tree derived from genomic data (Fig. 3a) placed strain F2^T^ and *Eubacterium pyruvativorans* KHGC13 within a strongly supported monophyletic clade, with 100% ultrafast bootstrap support This significantly exceeded the 61% support observed in the 16S rRNA gene tree, thereby confirming the close genomic relationship between these two strains. The clade containing strain F2^T^ and *Eubacterium pyruvativorans* KHGC13 was resolved as the immediate sister group of *Aminicella lysinilytoca* DSM 28287^T^, with 85% bootstrap support. The remaining *Aminicella* species formed a well-supported clade (97% bootstrap), within which *Aminicella nodata* ATCC 33099^T^ and *Aminicella minuta* ATCC 700079^T^ clustered together (100% bootstrap), and *Aminicella butyriciproducens* DSM 104962^T^ and *Aminicella sp000688015* AB3007 formed a separate clade (100% bootstrap). *Mogibacterium pumilum* ATCC 700696^T^ and *Eubacterium limosum* ATCC 8486^T^ were successively more distant outgroups, consistent with their classifications in distinct genera and families, respectively. The genome-based topology was fully congruent with the 16S rRNA gene tree at all major nodes, and the markedly improved bootstrap support throughout demonstrates the superior phylogenetic resolution afforded by multi-locus genome-based analysis compared to single-gene 16S rRNA phylogeny for genus-level delimitation.

**Fig. 3.**
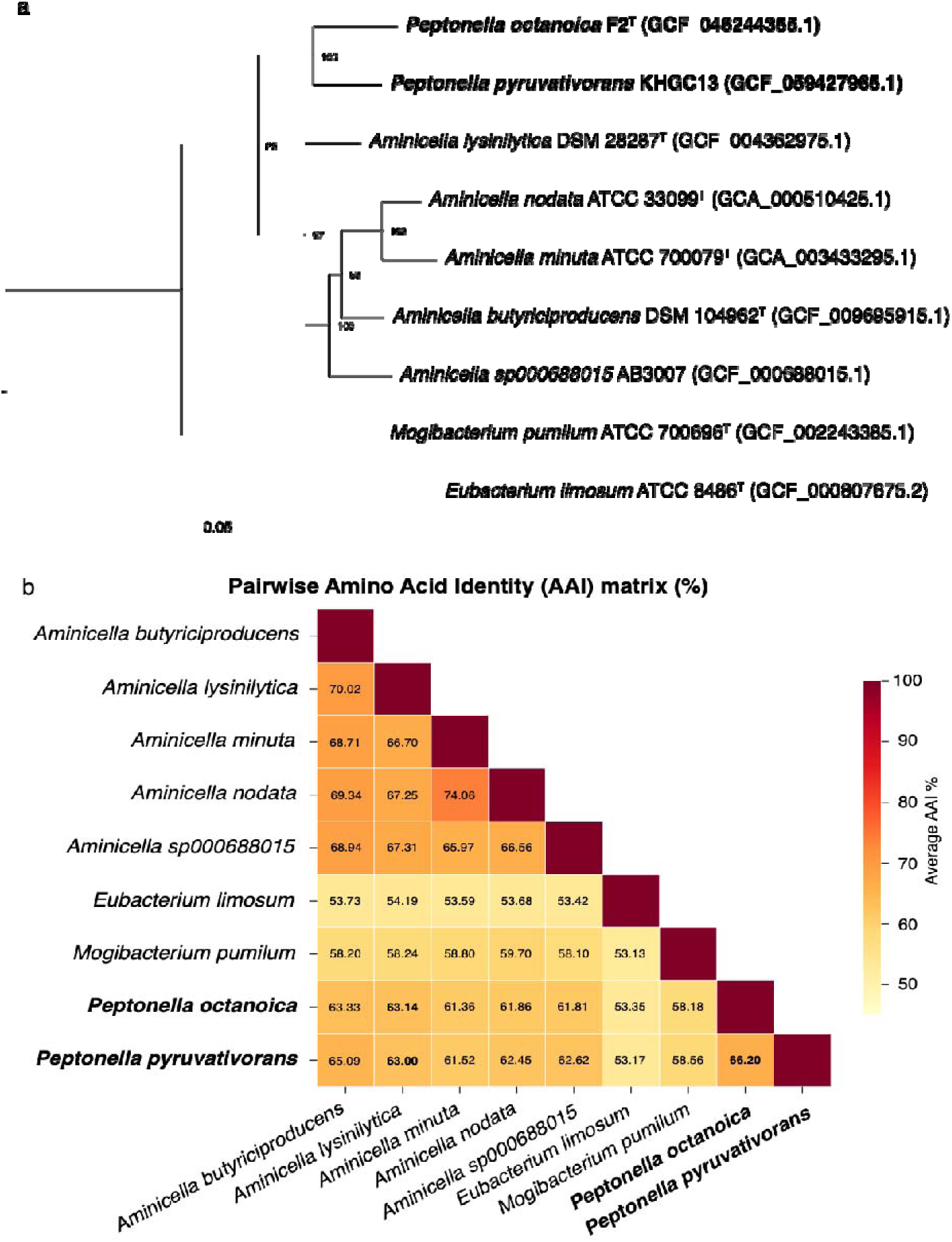
Phylogenomic characterization of *Peptonella octanoica* F2^T^ and *Peptonella pyruvativorans* KHGC13. (a) Maximum-likelihood phylogenetic tree inferred from a concatenated alignment of 120 conserved bacterial marker genes identified with GTDB-tk v2.7.2, utilizing the r232 reference database. Node numbers represent ultrafast bootstrap support values (%) from 1,000 replicates; only values of 50% or higher are displayed. Branch lengths are scaled in substitutions per site. *Eubacterium limosum* ATCC 8486^T^ was used as the outgroup. GenBank/RefSeq accession numbers of the genome sequences are given in parentheses. (b) Pairwise average amino acid identity (AAI) matrix computed with EzAAI v1.2.4. Values are expressed as percentages and represent the average of bidirectional comparisons. Cells are coloured according to AAI value, as indicated by the scale bar. Values highlighted in bold indicate pairwise comparisons involving the two Peptonella strains.

## Average amino acid identity

To further elucidate the genus– and species-level relationships of strain F2^T^ and *Eubacterium pyruvativorans* KHGC13 within a broader phylogenetic context, pairwise Average Amino Acid Identity (AAI) values were computed using EzAAI v1.2.4 [32]. Protein-coding sequences were predicted with Prodigal v2.6.3 [33], and pairwise comparisons were conducted utilizing MMseqs2 v17-b804f [34].

Pairwise AAI values across the nine genomes are shown in Figure 3b. *Peptonella octanoica* F2^T^ and *Peptonella pyruvativorans* KHGC13 exhibit 66.20% AAI, consistent with congeneric classification under the genus-level threshold of ≥65% proposed in the literature [35, 36]. The AAI values between strain F2^T^ and *Aminicella lysinilytica* DSM 28287^T^ (63.14%) and between KHGC13 and *A. lysinilytica* DSM 28287^T^ (63.00%) fall slightly below the 65% threshold but are markedly higher than those observed between the *Aminicella/Peptonella* clade and *Eubacterium limosum* ATCC 8486^T^ (53.17-54.19%), which belongs to a distinct family [24].

AAI values across all *Aminicella* species ranged from 65.97% to 74.06%, consistent with their classification within the same genus in the GTDB-tk database. In contrast, AAI values for *Mogibacterium pumilum* ATCC 700696^T^ (58.10-59.70%) were intermediate, reflecting its status as a related but distinct genus. The slight depression in AAI values among the *Peptonella* species indicates a divergent but closely related lineage within the same genus-level clade, as supported by phylogenomic and dDDH evidence.

## Discussion

The isolation of *Peptonella octanoica* strain F2^T^ from the SSO-fed chain-elongation reactor [15] holds considerable ecological and biotechnological importance. *P. octanoica* metabolizes lactate, acetate, and propanoate as substrates, yielding a mixture of short– and medium-chain carboxylates, with octanoate serving as the predominant MCCA when lactate is present. This substrate and product profile directly corresponds to the conditions and product spectrum observed in the reactor during periods of highest MCCA productivity. Notably, octanoate production accounted for up to 20% of total MCCAs in the bioreactor [15], reflecting a level of C8 specificity seldom observed in chain-elongation systems fed with complex organic waste streams [2, 3, 6]. We anticipate that the availability of *P. octanoica* strain F2^T^ as a pure-culture isolate will enable the elucidation of physiological principles governing C8 production.

Amplicon sequencing of the reactor microbiome identified *Eubacterium* (specifically, reads classified as *Eubacterium_T sp903789635* (GTDB r228), a taxon previously binned from metagenomic sequencing of a chain-elongation reactor [43]) as a dominant chain-elongating community member during periods of sustained MCCA production, consistent with strain F2^T^’s phylogenetic affinity for this lineage. Furthermore, the pH optimum of strain F2^T^ (pH 5-6) aligns with the reactor’s operating pH (pH 5.0) and is markedly lower than the near-neutral pH optima reported for most characterized CEB, including *Clostridium kluyveri* [10], indicating that strain F2^T^ is specifically adapted to the acidic conditions common in chain elongating bioreactors receiving food waste.

Phylogenomic and phenotypic evidence presented herein demonstrate that *P. octanoica* F2^T^ and *Eubacterium pyruvativorans* do not belong to *Eubacterium* sensu stricto but instead to a phylogenetically coherent clade, for which we propose the genus *Peptonella* gen. nov. This conclusion aligns with the long-recognized polyphyletic nature of the genus *Eubacterium* Prévot 1938, whose historically broad and taxonomically loose circumscription defined by exclusion rather than shared derived traits has resulted in the inclusion of phenotypically and phylogenetically diverse species. These species are distributed across multiple families within the phylum *Bacillota*, including *Eubacteriaceae, Lachnospiraceae, Eggerthellaceae,* and *Oscillospiraceae* [18]. The type species, *Eubacterium limosum*, belongs to a restricted cluster comprising only *E. limosum, E. barkeri,* and *E. callanderi*. It has been recommended that the genus name be restricted to this cluster, with all other species transferred to phylogenetically coherent genera [18]. The systematic disassembly of *Eubacterium* has been underway for thirty years, with previous members reassigned to *Agathobacter* [37], *Anaerobutyricum* [38], *Lachnoanaerobaculum* [39], *Pseudoramibacter* [24], *Peptoclostridium* [40], and *Aminicella* [41], among others.

*P. octanoica* and *E. pyruvativorans* share several key phenotypic characteristics consistent with their phylogenetic proximity, including strictly anaerobic growth, a monoderm cell wall ultrastructure, absence of spore formation, and the ability to utilize acetate and propanoate as substrates and to produce MCCAs via the reverse β-oxidation pathway [14, 47]. Additionally, *E. pyruvativorans* is a hyperammonia-producing (HAP) bacterium with an exceptionally high rate of ammonia production during amino acid fermentation (375 nmol mg protein-1 min-1) [42], a trait associated with its rumen habitat and ecological role in proteolysis; this property has also been observed in strain F2^T^. However, the two species can be clearly differentiated on several physiological and genomic properties. First, *E. pyruvativorans* was characterized primarily as an amino acid-fermenting organism that grows on pyruvate and trypticase as primary carbon and energy sources, with lactate supporting only weak growth [42], whereas strain F2^T^ grows efficiently on lactate as its primary electron donor. Second, *E. pyruvativorans* produces hexanoate as its dominant MCCA product [14], whereas strain F2^T^ produces octanoate as its predominant MCCA when lactate is available, indicating a two-carbon extension of the chain-elongation cycle relative to its closest relative. Third, the G+C content of strain F2^T^ (50.2 mol%) is lower than that of *E. pyruvativorans* KHGC13 (54.2 mol%), consistent with their status as distinct species as supported by dDDH.

*P. octanoica* also differs substantially from *Aminicella lysinilytica* DSM 28287^T^, the type species of the genus *Aminicella* and the closest relative of the *Peptonella* clade, in both 16S rRNA gene (90.12% similarity) and genome-based phylogenies. *A. lysinilytica* is a non-saccharolytic, strictly anaerobic Gram-positive rod isolated from a methanogenic reactor treating cattle farm waste. Its primary substrate is L-lysine, which it degrades in the presence of vitamin B12, producing acetate and butanoate as fermentation products. Importantly, no MCCA production has been reported in *A. lysinilytica* [41], and the capacity for chain elongation beyond butanoate appears unique to the *Peptonella* strains within the *Anaerovoracaceae* family. The G+C content of *A. lysinilytica* (45.5 mol%) differs substantially from that of the *Peptonella* strains. Also, the AAI value between *A. lysinilytica* and the *Peptonella* strains is 63-63.14%, which is beyond common genus thresholds (e.g., =>65%) [36].

Based on the phylogenomic, AAI, dDDH, and phenotypic evidence presented here, we propose the genus *Peptonella* gen. nov. to accommodate strain F2^T^ and *Eubacterium pyruvativorans*, with *Peptonella octanoica* sp. nov. designated as the type species and *Eubacterium pyruvativorans* Wallace et al. 2003 reclassified as *Peptonella pyruvativorans* comb. nov. The genus and species names have also been registered in the SeqCode registry (seqco.de/r:eu_6pd0e)[43]. The formal descriptions are provided below.

## DESCRIPTION OF *PEPTONELLA* GEN. NOV

*Peptonella* (Pep.to.nel’la – N.L. fem. n. *peptonum*, peptone; L. fem. dim. n. suff. –*ella*, diminutive ending; N.L. fem. n. *Peptonella*, the ability to use peptides in peptone-rich media).

Cells are strictly anaerobic, non-sporulating, produce medium-chain carboxylates, and possess a monoderm cell wall structure. The cells are non-saccharolytic. Phylogenetically, the genus represents a distinct lineage in the family *Anaerovoracaceae*. The type species is *Peptonella octanoica*.

## DESCRIPTION OF *PEPTONELLA OCTANOICA* SP. NOV

*Peptonella octanoica* (oc.ta.no’i.ca – N.L. neut. n. acidum *octanoicum*, octanoic acid; N.L. fem. adj. *octanoica*, referring to the production of octanoic acid)

Strictly anaerobic, Gram-stain-negative (but possessing a Gram-positive-type cell wall), non-spore-forming, rod-shaped bacterium. The average cell size is 1.5–2.5 × 0.27–0.33 µm, and the microorganism exhibits optimal growth in mDSM104 medium with lactate at 37 °C. Furthermore, cell growth occurs over a pH range of 5–8, with an optimum at pH 5–6. Fermentative end products include even-chain (acetate, butanoate, hexanoate, and octanoate), odd-chain (propanoate, pentanoate, and heptanoate), and branched-chain (iso-butanoate, iso-pentanoate, iso-hexanoate, and 5-methyl-hexanoate) carboxylates, as well as ammonium and gaseous products such as H_2_ and CO_2_. The cells utilize acetate and propanoate, elongating them into MCCAs in the presence of lactate. A complete gene cluster encoding the Rnf complex and the reverse β-oxidation pathway is present in the genome. Colonies are visible on mDSM104 agar plates after 3 days at 37 °C and appear white, small, circular, raised in elevation, and entire in margin after 5-7 days.

The type strain is F2^T^ (DSM=KCTC) and was isolated from an anaerobic bioreactor fermenting organic wastes. The DNA G+C content of the type strain is 50.2 mol%.

## DESCRIPTION OF PEPTONELLA PYRUVATIVORANS COMB. NOV

*Peptonella pyruvativorans* (py.ru.va.ti’vo.rans – N.L. neut. n. *pyruvatum*, pyruvate; L. pres. part. *vorans*, devouring, eating greedily; N.L. fem. part. adj. *pyruvativorans*, devouring pyruvate)

Basionym: *Eubacterium pyruvativorans* Wallace et al. 2003, Int J Syst Evol Microbiol 53:965–970.

The description is as given by Wallace et al. (2003) [14]. The type strain is I-6T (= ATCC BAA-574^T^ = NCIMB 13911^T^), isolated from the ruminal fluid of a sheep.

## Acknowledgements

We thank Jasmeen Parmar and Diana Dyussekenova for operating the bioreactor, and Ali Darbandi (Nanoscale Biomedical Imaging Facility, The Hospital for Sick Children Research Institute) for support with TEM analysis. We thank Konstantinos T. Konstantinidis and the curators of seqcode, Luis M. Rodriguez-R., Juan Valero Tebar and Marike Parmer, for their assistance with Latin naming of the organism.

## Author contributions

D.K.N. and B.G.L. performed investigations, data curation, formal analysis and validation. D.K.N. isolated the strain and wrote the original draft of the manuscript. D.K.N., B.G.L., and C.E.L. reviewed and edited the manuscript. C.E.L. conceptualized, supervised and acquired funding for this work.

## Data Availability

The GenBank and RefSeq accession numbers for the whole-genome sequence of strain F2^T^ are GCA_046244355.1 and GCF_046244355.1, respectively. Raw data have been deposited at the NCBI SRA database under the accession number SRR31952220. The accession number for the 16S rRNA gene sequence is PV739030.

