## Supplementary material for "*Peptonella octanoica* gen. nov., sp. nov., a new medium-chain carboxylate-producing bacterium, and the reclassification of *Eubacterium pyruvativorans* as *Peptonella pyruvativorans* comb. nov": Supp Info

### TYPE (STRAIN) GENOME SERVER

### JOB SUMMARY

| | |
|---|---|
| PRINT DATE: | 2026-07-02 21:35:48 +0200 |
| JOB ID: | cbdc8cbb-c23f-49f8-aadd-f81d03b71ab1 |
| RESULT PAGE: | https://tygs.dsmz.de/user_results/show?guid=cbdc8cbb-c23f-49f8-aadd-f81d03b71ab1 |

#### Table 1: Phylogenies

**Publication-ready versions** of both the genome-scale GBDP tree and the 16S rRNA gene sequence tree can be customized and exported either in SVG (vector graphic) or PNG format from within the phylogeny viewers in your TYGS result page. For publications the **SVG format is recommended** because it is lossless, always keeps its high resolution and can also be easily converted to other popular formats such as PDF or EPS. Please follow the link provided above!

#### Table 2: Identification

The below list contains the result of the TYGS species identification routine.

Explanation of remarks that might occur in the below table:

**remark [R1]:** The TYGS type strain database is automatically updated on an almost daily basis. However, if a particular type strain genome is not available in the TYGS database, this can have several reasons which are detailed in the FAQ. You can request an extended 16S rRNA gene analysis via the 16S tree viewer found in your result page to detect **not yet genome-sequenced** type strains relevant for your study.

**remark [R2]:** > 70% dDDH value (formula $d_4$) and (almost) minimal dDDH values for gene-content formulae $d_0$ and $d_6$ indicate a potentially unreliable identification result and should thus be checked via the 16S rRNA gene sequence similarity. Such strong deviations can, in principle, be caused by sequence contamination.

**remark [R3]:** G+C content difference of > 1 % indicates a potentially unreliable identification result because within species G+C content varies no more than 1 %, if computed from genome sequences (PMID: 24505073).

| Strain | Conclusion | Identification result | Remark |
|---|---|---|---|
| **'Aminicella sp905214225 (OCT)'** | potential new species | | see [R1] |

#### Table 3: Pairwise comparisons of user genomes vs. type-strain genomes

The following table contains the pairwise dDDH values between your user genomes and the selected type-strain genomes. The dDDH values are provided along with their confidence intervals (C.I.) for the three different GBDP formulas:

- formula $d_0$ (a.k.a. GGDC formula 1): length of all HSPs divided by total genome length
- formula $d_4$ (a.k.a. GGDC formula 2): sum of all identities found in HSPs divided by overall HSP length
- formula $d_6$ (a.k.a. GGDC formula 3): sum of all identities found in HSPs divided by total genome length

**Note:** Formula $d_4$ is independent of genome length and is thus robust against the use of incomplete draft genomes. For other reasons for preferring formula $d_4$, see the FAQ.

| Query | Subject | $d_0$ | C.I. $d_0$ | $d_4$ | C.I. $d_4$ | $d_6$ | C.I. $d_6$ | Diff. G+C Percent |
|---|---|---|---|---|---|---|---|---|
| **'Aminicella sp905214225 (OCT).fna'** | *Candidatus* Avitreponema avistercoris B3-4054 | 12.5 | [9.8 - 15.8] | 63.7 | [60.8 - 66.6] | 12.9 | [10.6 - 15.7] | 5.14 |
| **'Aminicella sp905214225 (OCT).fna'** | *Clostridium vitabionis* YH-T4B42 | 12.6 | [10.0 - 15.9] | 32.2 | [29.8 - 34.8] | 13.0 | [10.7 - 15.8] | 5.66 |
| **'Aminicella sp905214225 (OCT).fna'** | *Eubacterium nodatum* ATCC 33099 | 12.7 | [10.0 - 16.0] | 27.7 | [25.4 - 30.2] | 13.1 | [10.8 - 15.9] | 12.07 |
| **'Aminicella sp905214225 (OCT).fna'** | *Aminicella lysinilytica* DSM 28287 | 12.7 | [10.1 - 16.0] | 26.5 | [24.2 - 29.0] | 13.1 | [10.8 - 15.9] | 4.45 |
| **'Aminicella sp905214225 (OCT).fna'** | *Eubacterium minutum* ATCC 700079 | 12.8 | [10.1 - 16.0] | 26.4 | [24.1 - 28.9] | 13.1 | [10.8 - 15.9] | 4.41 |
| **'Aminicella sp905214225 (OCT).fna'** | *Eubacterium infirmum* NCTC 12940 | 12.7 | [10.0 - 16.0] | 26.3 | [24.0 - 28.8] | 13.1 | [10.8 - 15.9] | 10.0 |
| **'Aminicella sp905214225 (OCT).fna'** | *Clostridium fessum* SNUG30386T | 12.6 | [10.0 - 15.9] | 25.9 | [23.6 - 28.4] | 13.0 | [10.7 - 15.8] | 1.88 |
| **'Aminicella sp905214225 (OCT).fna'** | *Lachnospira hominis* CLA JM-H10 | 12.6 | [9.9 - 15.8] | 25.9 | [23.6 - 28.4] | 13.0 | [10.7 - 15.7] | 13.15 |
| **'Aminicella sp905214225 (OCT).fna'** | *Jutongia huaianensis* NSJ-37 | 12.6 | [9.9 - 15.9] | 24.6 | [22.3 - 27.1] | 13.0 | [10.7 - 15.7] | 6.12 |
| **'Aminicella sp905214225 (OCT).fna'** | *Senimuribacter intestinalis* DSM 106208T | 12.7 | [10.0 - 15.9] | 24.4 | [22.1 - 26.9] | 13.1 | [10.7 - 15.8] | 6.28 |
| **'Aminicella sp905214225 (OCT).fna'** | *Emergencia timonensis* SN18 | 12.6 | [9.9 - 15.9] | 24.1 | [21.8 - 26.6] | 13.0 | [10.7 - 15.8] | 4.39 |
| **'Aminicella sp905214225 (OCT).fna'** | *Treponema porcinum* ATCC BAA-908 | 12.5 | [9.9 - 15.8] | 23.8 | [21.5 - 26.3] | 12.9 | [10.6 - 15.7] | 7.72 |
| **'Aminicella sp905214225 (OCT).fna'** | *Pararoseburia lenta* NSJ-9 | 12.6 | [9.9 - 15.9] | 23.8 | [21.5 - 26.2] | 13.0 | [10.7 - 15.8] | 5.29 |
| **'Aminicella sp905214225 (OCT).fna'** | *Eubacterium sulci* ATCC 35585 | 12.7 | [10.0 - 16.0] | 23.7 | [21.4 - 26.2] | 13.1 | [10.7 - 15.8] | 10.3 |
| **'Aminicella sp905214225 (OCT).fna'** | *Allobaileyella intestinalis* DSM 106896T | 12.8 | [10.1 - 16.0] | 23.5 | [21.2 - 26.0] | 13.2 | [10.8 - 15.9] | 3.77 |
| **'Aminicella sp905214225 (OCT).fna'** | *Lactimicrobium massiliense* Marseille-P4301 | 12.6 | [10.0 - 15.9] | 23.1 | [20.8 - 25.6] | 13.0 | [10.7 - 15.8] | 2.92 |
| **'Aminicella sp905214225 (OCT).fna'** | *Hornefia porci* 68-3-10 | 12.9 | [10.2 - 16.2] | 22.6 | [20.3 - 25.1] | 13.3 | [10.9 - 16.0] | 2.12 |
| **'Aminicella sp905214225 (OCT).fna'** | *Hornefia butyriciproducens* DSM 104962T | 13.0 | [10.3 - 16.3] | 21.9 | [19.6 - 24.3] | 13.3 | [11.0 - 16.1] | 1.43 |
| **'Aminicella sp905214225 (OCT).fna'** | *Eubacterium pyruvativorans* I-6 | 13.4 | [10.6 - 16.7] | 21.6 | [19.3 - 24.0] | 13.7 | [11.3 - 16.5] | 4.67 |
| **'Aminicella sp905214225 (OCT).fna'** | *Eubacterium limosum* ATCC 8486 | 12.6 | [9.9 - 15.8] | 20.3 | [18.1 - 22.7] | 13.0 | [10.6 - 15.7] | 3.0 |

#### Table 4: Strains in your dataset

Joint dataset of automatically determined closest type strains (if this mode was chosen), manually selected type strains (if selected accordingly) and the provided user strains, if provided (marked in yellow).

| Strain | Authority | Other deposits | Synonyms | Base pairs | Percent G+C | No. proteins | Goldstamp | Bioproject accession | Biosample accession | Assembly accession | IMG OID |
|---|---|---|---|---|---|---|---|---|---|---|---|
| *Eubacterium minutum* ATCC 700079 | Poco et al. 1996 emend. Wade et al. 1999 | CIP 104795; DSM 18759; M-6 | *Eubacterium minutum* | 1870 942 | 45.8 | 1455 | Gp0386586 | PRJNA282954 | SAMN03897724 | GCA_003433295 | |
| *Hornefia porci* 68-3-10 | Wylensek et al. 2021 | DSM 104948; JCM 34388 | *Hornefia porci* | 2814 507 | 52.3 | 2548 | | PRJNA224116 | SAMN05726916 | GCF_001940235 | |
| *Jutongia huaianensis* NSJ-37 | Liu et al. 2022 | CGMCC 1.32810; KCTC 25089 | *Jutongia huaianensis* | 3060 005 | 44.1 | 2682 | | PRJNA656402 | SAMN15805243 | GCA_014384985 | |
| *Candidatus* Avitreponema avistercoris B3-4054 | Gilroy et al. 2021 | | *Candidatus* Avitreponema avistercoris | 1871 725 | 55.3 | 1772 | | PRJNA543206 | SAMN15816977 | GCA_017694555 | |
| *Lactimicrobium massiliense* Marseille-P4301 | Togo et al. 2019 | CSUR P4301 | *Lactimicrobium massiliense* | 2457 574 | 47.3 | 2346 | Gp0443118 | PRJNA224116 | SAMEA4587996 | GCF_900343155 | |
| *Clostridium vitabionis* YH-T4B42 | Shin et al. 2021 | KCTC 25105; NBRC 114767 | *Clostridium vitabionis* | 2858 048 | 55.8 | 2506 | | PRJNA224116 | SAMN16630968 | GCF_015351765 | |
| *Allobaileyella intestinalis* DSM 106896T | (Wylensek et al. 2021) Deshmukh et al. 2026 | JCM 34418; RF-744-FAT-WT-3 | *Allobaileyella intestinalis*; *Baileyella intestinalis* | 2014 858 | 46.4 | 1738 | | PRJNA224116 | SAMN12619154 | GCF_009695865 | |
| *Emergencia timonensis* SN18 | Bessis et al. 2016 | CSUR P2260 | *Emergencia timonensis* | 4659 729 | 45.8 | 4325 | | PRJEB13931 | SAMEA3959737 | GCA_900086585 | |
| *Clostridium fessum* SNUG30386T | Seo et al. 2021 | KCTC 15633; JCM 32258 | *Clostridium fessum* | 3259 355 | 48.3 | 2839 | | PRJNA438872 | SAMN08731216 | GCA_003024715 | |
| *Eubacterium infirmum* NCTC 12940 | Cheeseman et al. 1996 | ATCC 700433; W 1471 | *Eubacterium infirmum* | 1906 995 | 40.2 | 1745 | Gp0384418 | PRJEB6403 | SAMEA48403918 | GCA_900450455 | |

| Strain | Authority | Other deposits | Synonyms | Base pairs | Percent G+C | No. proteins | Goldstamp | Bioproject accession | Biosample accession | Assembly accession | IMG OID |
|---|---|---|---|---|---|---|---|---|---|---|---|
| *Aminicella lysinilytica* DSM 28287 | Ueki et al. 2015 | JCM 19863; WN037 | *Aminicella lysinilytica* | 2282262 | 45.7 | 2123 | Gp0325700 | PRJNA519323 | SAMN10872748 | GCA_004362975 | 2795385453 |
| *Treponema porcinum* ATCC BAA-908 | Nordhoff et al. 2005 emend. Hördt et al. 2020 | 14V28; CIP 108245; JCM 12342 | *Treponema porcinum* | 2507489 | 42.5 | 2192 | Gp0088921 | PRJNA245589 | SAMN02745149 | GCA_900167145 | 2582581326 |
| *Eubacterium nodatum* ATCC 33099 | Holdeman et al. 1980 | CIP 104213; CCUG 15996; DSM 3993; JCM 14550; JCM 9977; VPI D6A-5 | *Eubacterium nodatum* | 1829558 | 38.1 | 1691 | Gp0004475 | PRJNA89653 | SAMN00829151 | GCA_000510425 | 2558860175 |
| *Eubacterium sulci* ATCC 35585 | (Cato et al. 1985) Jalava and Eerola 1999 | CCUG 20560; VPI D45A-29A | *Eubacterium sulci*; *Fusobacterium sulci* | 1728669 | 39.9 | 1587 | Gp0005400 | PRJNA53047 | SAMN02641587 | GCA_000526055 | 2558860286 |
| *Eubacterium limosum* ATCC 8486 | (Eggerth 1935) Prévot 1938 | CIP 104169; CCUG 16793; DSM 20543; JCM 6421; JCM 9978 | *Bacteroides limosus*; *Butyribacterium limosum*; *Eubacterium limosum* | 4369935 | 47.2 | 4084 | Gp0117762 | PRJNA270275 | SAMN03265381 | GCA_000807675 | |
| *Eubacterium pyruvativorans* I-6 | Wallace et al. 2003 | NCIMB 13911; ATCC BAA-574 | *Eubacterium pyruvativorans* | 2158512 | 54.8 | 1898 | Gp0112433 | PRJEB15823 | SAMN04487889 | GCA_900102225 | |
| *Senimuribacter intestinalis* DSM 106208T | Afrizal et al. 2025 | JCM 37188; YCFAG-7-CC-SB-Schm-I | *Senimuribacter intestinalis* | 3225420 | 43.9 | 3093 | | PRJEB50452 | SAMEA14418523 | GCA_943192975 | |
| *Lachnospira hominis* CLA JM-H10 | Hitch et al. 2025 | DSM 114599; LMG 33585 | *Lachnospira hominis* | 3148779 | 37.0 | 2947 | | PRJNA996881 | SAMN40466799 | GCA_040096395 | |
| *Pararoseburia lenta* NSJ-9 | Abdugheni et al. 2022 | CGMCC 1.32469; KCTC 15957 | *Pararoseburia lenta*; *Roseburia lenta* | 2552251 | 44.9 | 2314 | | PRJNA656402 | SAMN15808071 | GCA_014287435 | |

| Strain | Authority | Other deposits | Synonyms | Base pairs | Percent G+C | No. proteins | Goldstamp | Bioproject accession | Biosample accession | Assembly accession | IMG OID |
|---|---|---|---|---|---|---|---|---|---|---|---|
| *Hornefia butyriciproducens* DSM 104962T | Wylensek et al. 2021 | JCM 34390; WCA-MUC-591-APC-3H | *Hornefia butyriciproducens* | 2436683 | 51.6 | 2191 | | PRJNA224116 | SAMN12619152 | GCF_009695915 | |
| Aminicella sp905214225 (OCT).fna | | | | 2425940 | 50.2 | 2091 | | | | | |

#### Methods, Results and References

The genome sequence data were uploaded to the Type (Strain) Genome Server (TYGS), a free bioinformatics platform available under https://tygs.dsmz.de, for a whole genome-based taxonomic analysis [1]. The analysis also made use of recently introduced methodological updates and features [2,3]. Information on nomenclature, synonymy and associated taxonomic literature was provided by TYGS's sister database, the List of Prokaryotic names with Standing in Nomenclature (LPSN, available at https://lpsn.dsmz.de) [2,3]. The results were provided by the TYGS on 2026-06-23. The TYGS analysis was subdivided into the following steps:

##### Determination of closely related type strains

The determination of closely related type strains did not succeed because not a single 16S rDNA gene sequence was detected in the provided user genomes. The subsequent analyses are thus only based on the provided genome data and the manually selected type strains, if any.

##### Pairwise comparison of genome sequences

For the phylogenomic inference, all pairwise comparisons among the set of genomes were conducted using GBDP and accurate intergenomic distances inferred under the algorithm 'trimming' and distance formula $d_5$ [4]. 100 distance replicates were calculated each. Digital DDH values and confidence intervals were calculated using the recommended settings of the GGDC 4.0 [2,4].

##### Phylogenetic inference

The resulting intergenomic distances were used to infer a balanced minimum evolution tree with branch support via FASTME 2.1.6.1 including SPR postprocessing [5]. Branch support was inferred from 100 pseudo-bootstrap replicates each. The trees were rooted at the midpoint [6] and visualized with PhyD3 [7].

##### Type-based species and subspecies clustering

The type-based species clustering using a 70% dDDH radius around each of the 20 type strains was done as previously described [1]. The resulting groups are shown in Table 1 and 4. Subspecies clustering was done using a 79% dDDH threshold as previously introduced [8].

#### Results

##### Type-based species and subspecies clustering

The resulting species and subspecies clusters are listed in Table 4, whereas the taxonomic identification of the query strains is found in Table 1. Briefly, the clustering yielded 20 species clusters and the provided query strains were assigned to 1 of these. Moreover, user strains were located in 1 of 20 subspecies clusters.

##### Figure caption genome tree

**Figure 2.** Tree inferred with FastME 2.1.6.1 [5] from GBDP distances calculated from genome sequences. The branch lengths are scaled in terms of GBDP distance formula $d_5$. The numbers above branches are GBDP pseudo-bootstrap support values > 60 % from 100 replications, with an average branch support of 16.2 %. The tree was rooted at the midpoint [6].
